# Bacterial secondary metabolite biosynthetic potential in soil varies with phylum, depth, and vegetation type

**DOI:** 10.1101/818815

**Authors:** Allison M. Sharrar, Alexander Crits-Christoph, Raphaël Méheust, Spencer Diamond, Evan P. Starr, Jillian F. Banfield

**Affiliations:** Department of Earth and Planetary Science, University of California, Berkeley, CA, USA; Department of Plant and Microbial Biology, University of California, Berkeley, CA, USA; Innovative Genomics Institute, Berkeley, CA, USA

## Abstract

Bacteria isolated from soils are major sources of specialized metabolites, including antibiotics and other compounds with clinical value that likely shape interactions among microbial community members and impact biogeochemical cycles. Yet, isolated lineages represent a small fraction of all soil bacterial diversity. It remains unclear how the production of specialized metabolites varies across the phylogenetic diversity of bacterial species in soils, and whether the genetic potential for production of these metabolites differs with soil type. We sampled soils and saprolite from three sites in a northern California Critical Zone Observatory with varying vegetation and bedrock characteristics and used metagenomic sequencing and assembly to reconstruct 1,334 microbial genomes containing diverse biosynthetic gene clusters (BGCs) for secondary metabolite production. We obtained genomes for prolific producers of secondary metabolites, including novel groups within the Actinobacteria, Chloroflexi and candidate phylum Dormibactereota. Surprisingly, one genome of a Candidate Phyla Radiation bacterium encoded for a ribosomally synthesized linear azole/azoline-containing peptide, a capacity we found in other publicly available CPR bacterial genomes. Overall, bacteria with higher biosynthetic potential were enriched in shallow soils and grassland soils, with patterns of abundance of BGC type varying by taxonomy.

## Introduction

Many soil microbes biosynthesize secondary metabolite molecules that play important ecological roles in their complex and heterogeneous microenvironments. Secondary (or “specialized”) metabolites are auxiliary compounds that microbes produce which are not required for normal cell growth, but which benefit the cells in other ways. These compounds can have roles in nutrient acquisition, communication, inhibition, or other interactions with surrounding organisms or the environment^1^. Examples of these molecules include antibiotics^1^, siderophores^2^, quorum-sensing molecules^3^, immunosuppressants^4^, and degradative enzymes^5^.

Secondary metabolites are of interest for both their ecological and biogeochemical effects, as well as their potential for use in medicine and biotechnology. Antibiotics are a class of secondary metabolites with obvious human importance. Historically, antibiotic discovery relied on being able to culture organisms from the environment; however, the vast majority of environmental taxa cannot be cultured using current methods. Most known antibiotics are from cultured members of Actinobacteria, Proteobacteria, and Firmicutes^6^. Because soil microbial communities are so diverse and most microbial taxa in soil have not been well-described^7^, they offer a wealth of potential for the discovery of new and important microbial products.

Secondary metabolites are produced by biosynthetic gene clusters (BGCs), groups of colocated genes that function together to build a molecule. Nonribosomal peptide synthetases (NRPSs) and polyketide synthases (PKSs) are two of the largest classes of BGCs, encompassing most known antibiotics and antifungals^6^. NRPSs are characterized by condensation (CD) and adenylation (AD) domains^8^ and PKSs contain ketosynthase (KS) domains and a variety of other enzymatic domains^9^. These characteristic domains can be used to identify novel NRPS and PKS gene clusters and their abundances can be used as a proxy for biosynthetic potential^10^.

Little is known about how environmental variables impact the distribution of secondary metabolites in soil. Recent studies of microbial biosynthetic potential in soil have utilized amplicon sequencing of NRPS and PKS domains^11,12,13,14,15^. One study involving soils from a variety of environments demonstrated that NRPS and PKS domain richness was high in arid soils and low in forested soils^11^. Others showed that the composition of these domains correlated with latitude^12^ and vegetation^13^ at a continental scale and was distinct between urban and nonurban soils^14^. Because these studies rely on degenerate PCR primers designed for known domains, only sequences similar to known domains can be recovered. In contrast, genome-resolved metagenomics is able to recover divergent sequences within their genomic and phylogenetic context. Recently, this approach revealed abundant biosynthetic loci in Acidobacteria, Verrucomicrobia, Gemmatimonadetes, and the candidate phylum Rokubacteria^16^.

Here, we reconstructed genomes from 129 metagenomes sampled from various soil types and soil conditions in three northern California ecosystems and searched them for BGCs. The first site is a meadow grassland, the second is a Douglas fir / Madrone forested hillslope, and the third is an Oak grassland hillslope. A subset of the genomes from meadow grassland soils analyzed here were previously reported in studies that both reported on novel BGCs^16^ and demonstrated that soil depth and soil moisture affect microbial community structure and function^17,18^. We present a comparative analysis of the biosynthetic potential of bacteria from many phyla and report how soil microbiology and secondary metabolic potential vary with soil type and environmental conditions. Metagenomic studies such as this have the ability to identify new environmental and taxonomic targets key to the understanding of secondary metabolism ecology, and for the development of microbial natural products of human interest.

## Materials & Methods

### Sampling sites

Soil and saprolite samples were taken in areas studied by the Eel River Critical Zone Observatory (CZO). Samples were taken from soil depths of 20-200 cm over a four year period, from 2013 to 2016. The Eel River CZO experiences a Mediterranean climate characterized by hot, dry summers and cool, wet winters. The first fall rain after the dry summer generally comes in mid-late September and most rain falls between November and March. Average yearly rainfall varies from about 1.8-2 m^19^.

Samples were collected at two sites within the Angelo Coast Range Reserve: Rivendell, a forested hillslope^20,21^, and a nearby meadow^17,18,22^. The meadow and Rivendell are 1.5 km apart (Fig. 1). Both are underlain by the Coastal Belt of the Franciscan Formation, which consists of mostly argillite (shale), with some sandstone and conglomerate^23^. At Rivendell, the soil mostly lacks distinct horizons^24^ and varies in depth from 30-75 cm with saprolite directly below^19^. The northern slope of Rivendell is dominated by Douglas fir (*Pseudotsuga menziesii*) trees, while the southern slope has more Pacific madrone (*Arbutus menziesii*) trees. In the Angelo meadow, grass roots are confined to depths of <10 cm^17^.

**Figure 1.**
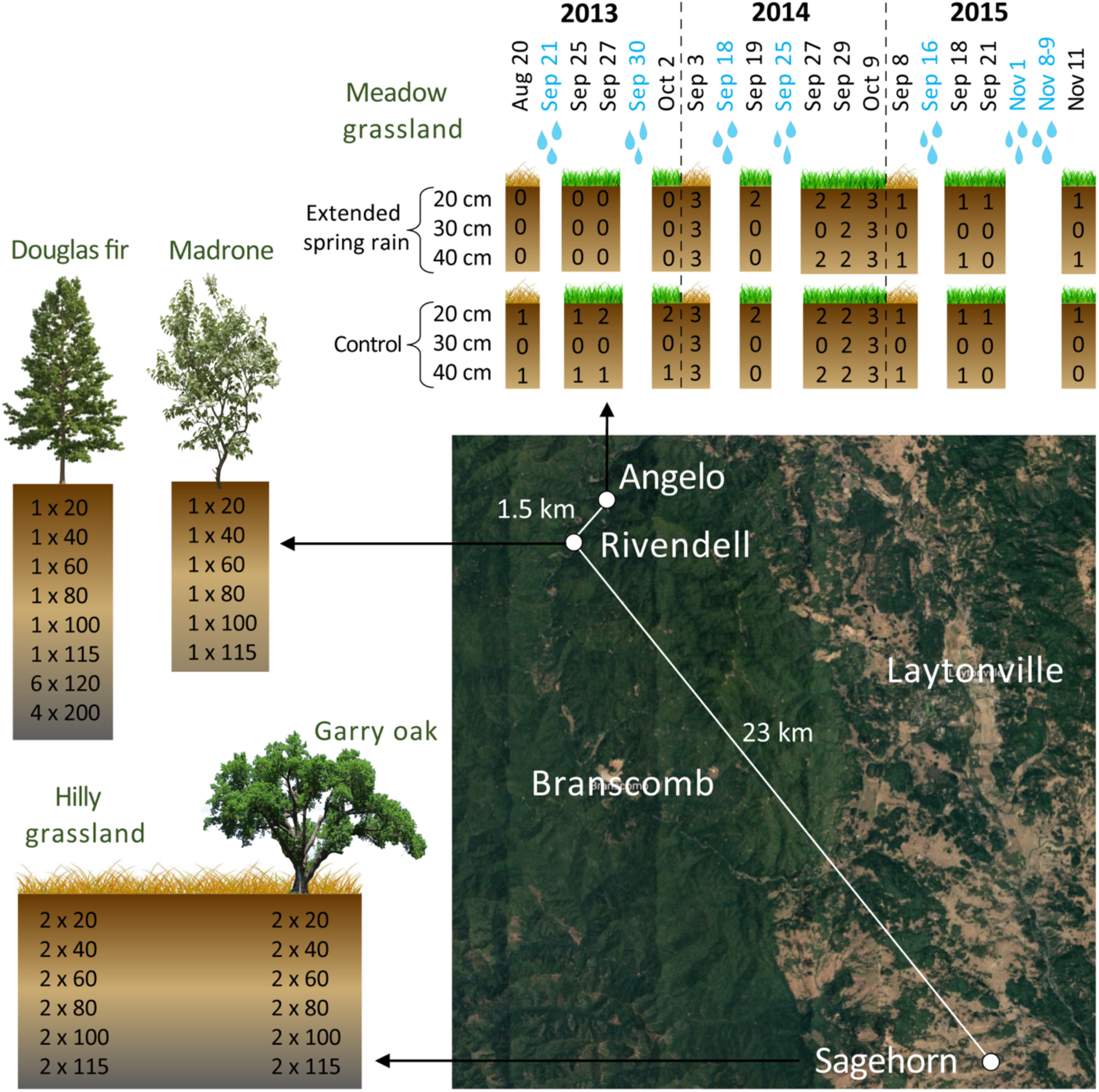
Eel River CZO sampling scheme. Soil and saprolite samples were taken from depths of 20-200 cm across three sites between 2013 and 2016. At **Angelo**, meadow grassland samples were taken before and after the first fall rains in 2013-2015 on the dates shown (blue = natural rainfall events). Numbers in the boxes show how many samples were taken at each depth on each date from either control plots or plots with experimentally extended spring rainfall. At **Rivendell**, samples were taken from both the north slope, under a Douglas fir tree, and the south slope, under a Madrone tree. Numbers in the boxes show how many samples were taken x depth (cm). Similarly at **Sagehorn**, samples were taken from below a Garry oak tree and in the nearby hilly grassland.

A third study site, Sagehorn, is a hilly grassland located about 23 km to the southeast of the other two sites (Fig. 1)^25,26^. Sagehorn is underlain by the Central Belt of the Franciscan Formation, a mélange with a sheared argillaceous matrix containing blocks of sandstone and other lithologies^27^. Sagehorn soils generally have a 30 cm thick organic-rich mineral A horizon underlain by a 10-20 cm thick Bt horizon with a high clay content, directly above saprolite. The low porosity mélange bedrock causes these layers to become entirely saturated in the winter wet season^26^. Sagehorn is primarily a grassland with scattered Garry oak (*Quercus garryana*) trees.

### Sampling and DNA extraction

At the meadow sites, 10 samples were taken on four dates in 2013 spanning before and after the first two fall rain events at soil depths of either 20 or 40 cm, as described in a previous publication^17^. In 2014, 60 samples were taken on five dates before and after the first two fall rain events at soil depths of 20, 30, and 40 cm. Samples came from six different plots: three treatment plot replicates with artificially extended spring rain and three control plot replicates, as described in Diamond et al^18^. In 2015, 13 samples were taken on four dates spanning before and after the first few fall rain events at soil depths of either 20 or 40 cm on either a control or treatment plot (Fig. 1). All Angelo samples will be referred to as the ‘meadow grassland’ samples.

At Rivendell, a depth profile of six samples (20, 40, 60, 80, 100, and 115 cm) was taken on the Douglas fir-dominated northern slope in 2013. Sterile scoops were used to sample soil and saprolite from a bucket auger. Samples were scooped directly into sterile Whirl-Pak bags and flash frozen onsite in dry ice and ethanol. In 2015, 10 deep saprolite samples were taken from the northern slope. A trackhoe outfitted with a coring auger was used to drill into the hillslope saprolite beneath mature Douglas fir trees. At depths of 120 and 200 cm, samples were taken using a sterilized hand auger. All samples from the northern slope of Rivendell will be referred to as the ‘Douglas fir’ samples. In 2016, a similar depth profile of six samples (20-115 cm) was taken on the southern slope under a Pacific madrone tree (the ‘Madrone’ samples). A soil pit was dug using a jackhammer and the wall of the pit was sampled with sterile scoops into 50 mL Falcon tubes which were immediately flash frozen on dry ice.

At Sagehorn, a depth profile of 12 samples (two at each depth: 20, 40, 60, 80, 100, and 115 cm) was taken from under a Garry oak tree (the ‘Garry oak’ samples) and from the nearby grassland (the ‘hilly grassland’ samples), for a total of 24 samples. The two soil pits were dug using a jackhammer. The walls of the pits were sampled on both sides with a sterile scoop, resulting in two samples per soil depth approximately 10 cm apart laterally. Samples were scooped into sterile 50 mL Falcon tubes which were immediately flash frozen on dry ice.

All samples were transported on dry ice and stored at -80°C until DNA extraction. In all cases, DNA was extracted from 10 g of material with the MoBio Laboratories PowerMax Soil DNA Isolation kit, using a previously described protocol^17^. This resulted in a total of 129 metagenomic samples across the Eel River CZO (Table S1).

### DNA sequencing, assembly, and genome reconstruction

All metagenomic library preparation and DNA sequencing was done at the Joint Genome Institute. 2013 Douglas fir samples and 2013-2014 meadow grassland samples were sequenced using 250 base pair (bp) paired-end Illumina reads. 2014 meadow grassland reads were quality-trimmed to 200 bp and assembled into individual metagenomes using a combination of IDBA-UD^28^ and MEGAHIT^29^, as previously described^18^. All other metagenomes were sequenced using 150 bp paired-end Illumina reads and datasets individually assembled using IDBA-UD^28^. Open reading frames were predicted with Prodigal^30^ and annotated by using USEARCH^31^ to search for similarity against UniProt^32^, UniRef90, and KEGG^33^ databases.

This dataset includes genomes binned from prior studies^17,18^ and newly reported genomes. Newly reported genomes from 2015 meadow grassland samples were binned using differential coverage binners ABAWACA2^34^, MaxBin2^35^, CONCOCT^36^, and MetaBAT^37^. Scaffolds from all other metagenomes were binned using ABAWACA2^34^, MaxBin2^35^, and MetaBAT^37^. For all metagenomes binned with multiple automated binners, the highest quality bins from each metagenome were selected using DasTool^38^.

### Ribosomal protein S3 (rpS3) analysis

All predicted proteins from the 129 metagenomes were searched for ribosomal protein S3 (rpS3) sequences using a custom Hidden Markov Model (HMM) from Diamond et al^18^. Only rpS3 proteins with lengths in the 120-450 amino acid range were considered, resulting in 20,789 rpS3 proteins. RpS3 protein taxonomy was identified at the phylum level using USEARCH^31^ to search against a database of rpS3 proteins from Hug et al^39^. RpS3 proteins were then clustered at 99% amino acid identity using USEARCH. This resulted in 7,013 de-replicated rpS3 sequences, each representing an approximately species-level cluster. Reads from each sample were mapped against the largest rpS3-containing scaffold in each cluster using Bowtie2^40^. Read mappings were filtered for ≥98% sequence identity and a coverage table was created by calculating coverage per base pair. The coverage table was normalized for sample sequencing depth using the formula: (coverage / reads input to the sample’s assembly) x average number of reads input to assemblies. The coverage table was used as input to a principal coordinate analysis (PCoA). First, a distance matrix was created in Python using the SciPy module scipy.spatial.distance.pdist^41^ with the Bray-Curtis distance metric. Then the PCoA was performed using the Scikit-Bio module skbio.stats.ordination.pcoa^42^.

### Genome filtering and de-replication

Bins were initially filtered for completeness and contamination based on the inventory of of 38 archaeal single copy genes or 51 bacterial single copy genes, except for CPR bacteria, where a reduced set of 43 CPR-specific genes was used^43^. Bins that had at least 70% of the single copy genes in their respective set with <4 having multiple copies were kept in the analysis. Next, CheckM^44^ lineage_wf was run on these bins, with a threshold of >70% complete with <10% contamination (for non-CPR bins only). To achieve the final draft genome set, bins were de-replicated at 98% nucleotide identity using dRep^45^ (Olm et al., 2017).

### Tree building and taxonomic determination

Genomes with >50% of their genes annotated to have best hits in one phylum were automatically assigned to that phylum. To check this phylum classification and identify the remaining genomes, a maximum-likelihood tree was calculated based on the concatenation of 16 ribosomal proteins (L2, L3, L4, L5, L6, L14, L15, L16, L18, L22, L24, S3, S8, S10, S17, and S19). Sequences were aligned using MAFFT^46^ version 7.390 (--auto option). Each alignment was further trimmed using trimAl^47^ version 1.4.22 (--gappyout option) before being concatenated. Tree reconstruction was performed using IQ-TREE^48^ version 1.6.6 (as implemented on the CIPRES web server^49^), using ModelFinder^50^ to select the best model of evolution (LG+F+I+G4), and with 1000 ultrafast bootstrap replicates. Using the same method, but with model LG+I+G4, a tree was created to compare Actinobacteria in this study to NCBI references from each Actinobacteria genus. All trees were visualized with iTol^51^.

### Differential abundance analysis

Reads from all 129 metagenomes were mapped against the de-replicated set of genomes using Bowtie2^40^. Raw read counts for each genome across each sample were input into DESeq2^52^ using R. Differential abundance across depth, controlling for site, was tested using the the DESeq2 model: design = ∼ site + depth. Each genome with a p-value adjusted for false discovery rate (FDR) of <0.05 for either increasing with depth (deep enriched) or decreasing with depth (shallow enriched) was put into the respective category. Differential abundance with vegetation was tested using the DESeq2 model: design = ∼ site + vegetation. Vegetation was classified simply as either grassland (meadow and hilly grassland samples) or tree-covered (Douglas fir, Madrone, and Garry oak samples). Each genome with an FDR adjusted p-value of <0.05 for either tree-covered or grassland enriched was put into the respective category.

### Biosynthetic gene cluster (BGC) analysis

To identify biosynthetic gene clusters (BGCs), antiSMASH 4.0^53^ was run on the final de-replicated set of genomes using default parameters. Only BGCs on contigs >10 kb were considered. Ketosynthase (KS) and condensation (CD) domains were identified using Pfam^54^ HMMs PF00109 and PF00668, respectively. BGC type was determined from the antiSMASH output. Only classes with >1% overall abundance were named in figures, others were put into the ‘Other’ category. This category is made up of both <1% abundant classes and BGCs that antiSMASH could not confidently place into a type category. The Wilcoxon signed-rank test was used in R to calculate significant differences between the average numbers of BGCs, KS and CD domains per genome between groups of genomes showing enrichment with depth or vegetation.

## Results

### Microbial community structure across the Eel River CZO

To compare microbial community composition across samples with varying bedrock lithology, vegetation, depth, and time since the last rainfall, assembled sequences of ribosomal protein S3 (rpS3) were used as marker genes for identifying different taxa. RpS3 is a universal single copy gene, assembles well from metagenomic data, and is recovered more frequently than whole genomes, which allows for a more inclusive view of microbial communities^55^. Our RpS3 analysis indicates that microbial communities across the Eel River CZO are generally dominated by the same bacterial phyla (Proteobacteria, Acidobacteria, Actinobacteria, Verrucomicrobia, Chloroflexi, and Gemmatimonadetes), but community composition is distinct between sampling sites and at different depths within the same site (Fig. 2A). Archaea are abundant, making up as much as 30% of the community in some samples. Some CPR bacteria are present at low abundance (usually <1% of the community) in most samples except in the meadow grassland samples. Acidobacteria are very abundant in Douglas fir and Madrone soil, whereas Actinobacteria are very abundant in the hilly grassland soil, especially at depth. At the meadow grassland, Archaea, Rokubacteria, and Nitrospirae are more abundant with depth.

**Figure 2.**
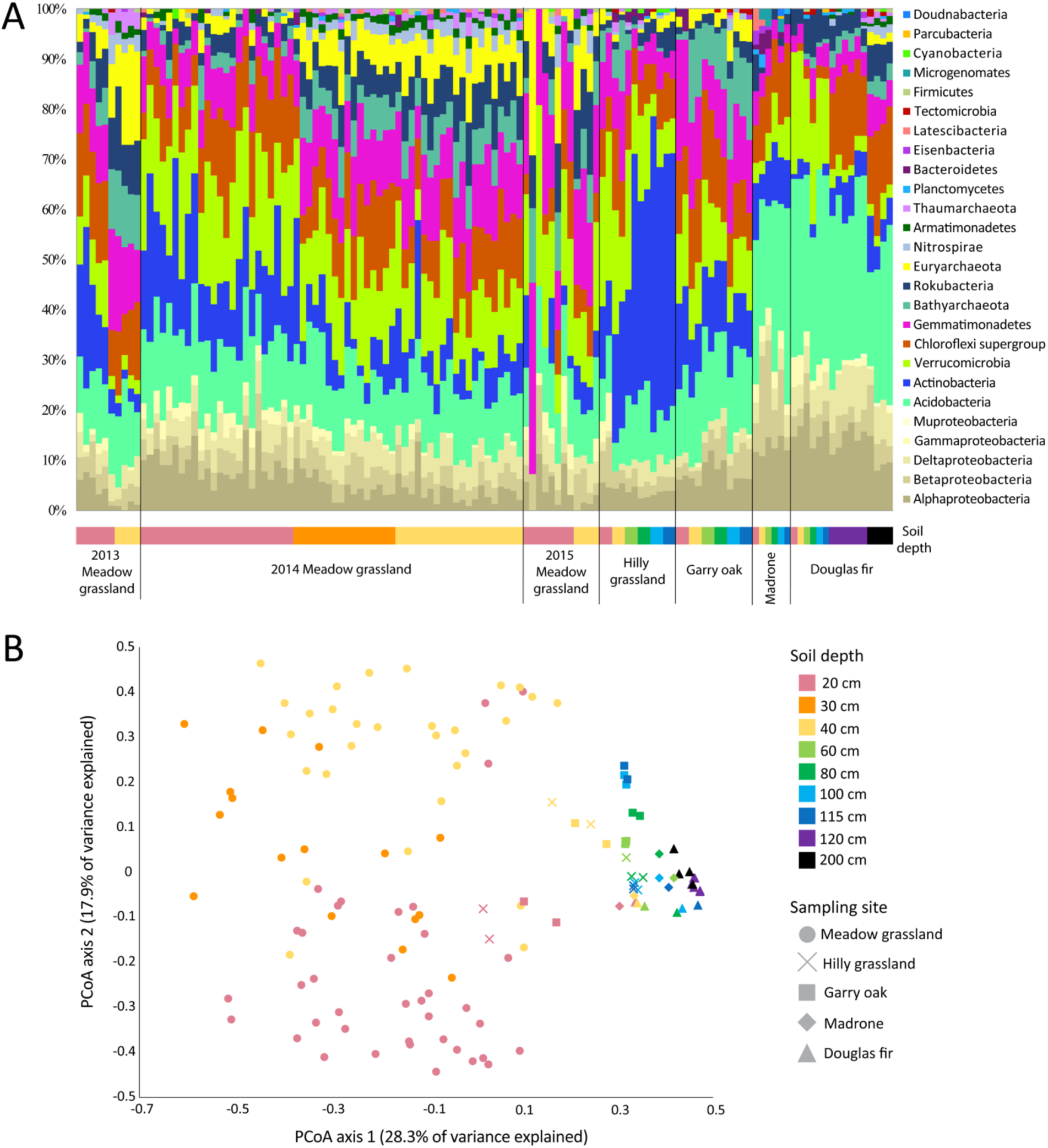
Microbial community structure across the Eel River CZO. **A**, Relative abundance of microbial phyla making up >1% of the community based on coverage of ribosomal protein S3 (rpS3)-containing contigs across the Eel River CZO metagenomes (*n* = 129). Samples are grouped by site and either year sampled or vegetation (black vertical lines), then by soil depth (colored bar below with same legend as Fig. 2B). **B**, Principal coordinates analysis (PCoA) of microbial community composition derived from read mapping of de-replicated rpS3-containing contigs and Bray-Curtis dissimilarities. Each point is one metagenomic sample (*n* = 129).

Of the varying environmental characteristics considered, sampling site and depth appear to be the greatest predictors of microbial community similarity (Fig. 2B). Communities separate by sampling site along the 1st PCoA axis (28.3% of variance explained). Meadow grassland samples then separate by depth along PCoA axis 2 (17.9% of variance explained). The meadow grassland and Douglas fir sites appear to have the most dissimilar microbial communities. Shallow (20 and 40 cm) hilly grassland and Garry oak samples begin to cluster with meadow grassland samples according to depth.

### Genome recovery and biosynthetic potential across taxonomic groups

We reconstructed 15,473 genomic bins from the 129 metagenomes. This set was narrowed to 3,895 bins after consideration of genome completeness and contamination, and dereplicated to a final set of 1,334 genomic bins used in subsequent analyses (Table S2), 944 of which were previously unpublished.

Overall, 3,175 BGCs were identified on contigs >10 kb across the 1,334 de-replicated genomes (Table S3). These genomes belonged to 22 different bacterial phylum-level groups, most of which showed some level of biosynthetic potential, and three archaeal phyla (Fig. 3; Fig. S1). Bacteria from certain phyla, such as the candidate phylum Rokubacteria, consistently have moderate numbers of BGCs in their genomes (Fig. 4A). Other phyla, such as Actinobacteria and Chloroflexi, have lower median values, but contain individual genomes with exceptionally high numbers of BGCs (Fig. 4A). Average amounts of KS and CD domains per genome also vary by phylum (Fig. 4B-C). Rokubacteria commonly have moderate amounts of KS domains but rarely many CD domains, whereas Acidobacteria more often have high amounts of CD domains.

**Figure 3.**
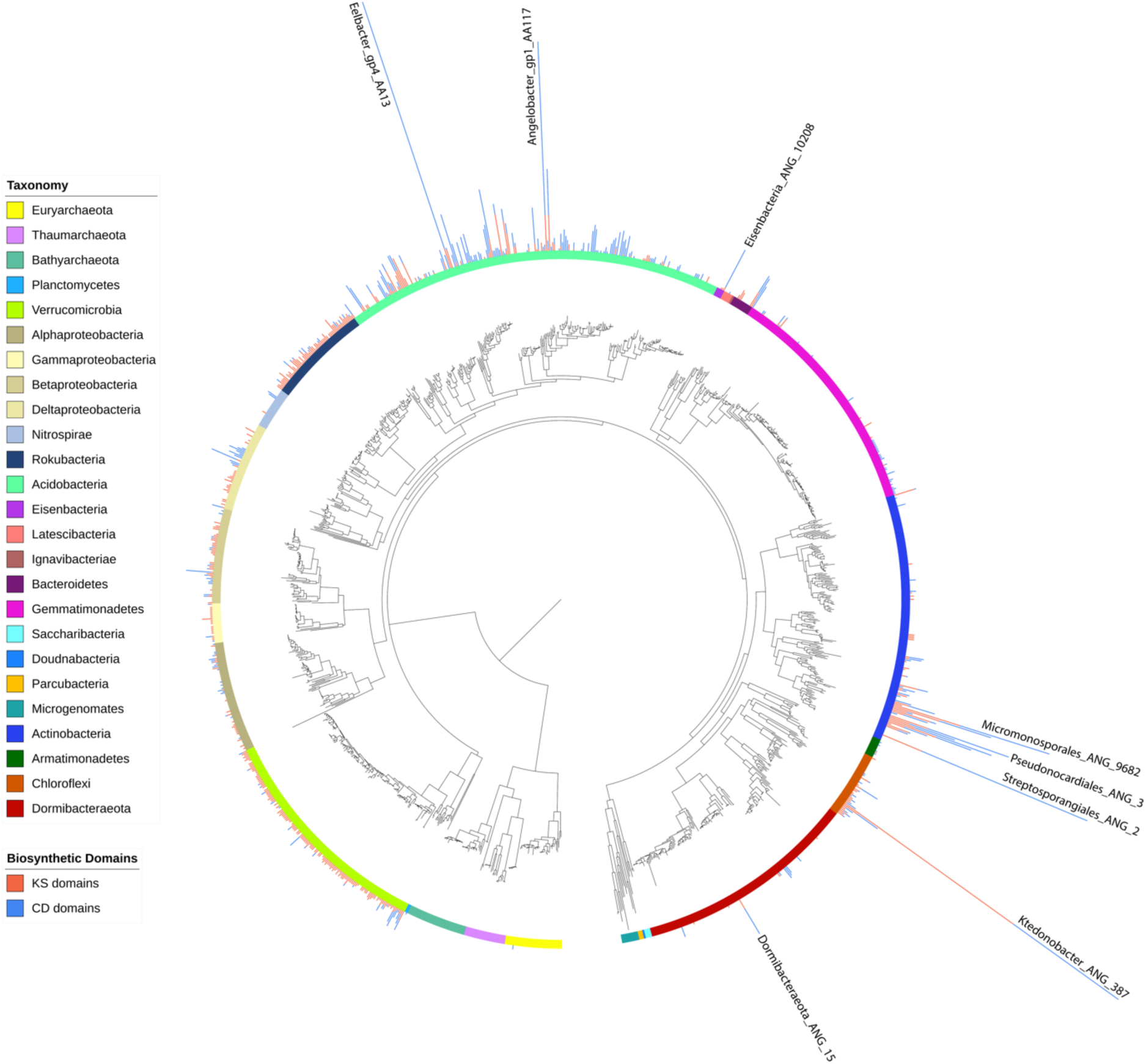
Concatenated ribosomal protein tree of high quality, de-replicated genomes. Maximum-likelihood tree based on the concatenation of 16 ribosomal proteins from genomes from all 129 metagenomic samples that passed thresholds of >70% complete and <10% contamination, according to CheckM (*n* = 1,334). Colored ring indicates taxonomic group. Stacked bar plots show the amount of KS (red) and CD (blue) domains identified by antiSMASH in each genome. Figure S1 is a higher resolution version of this figure with genome names and bootstrap information included.

**Figure 4.**
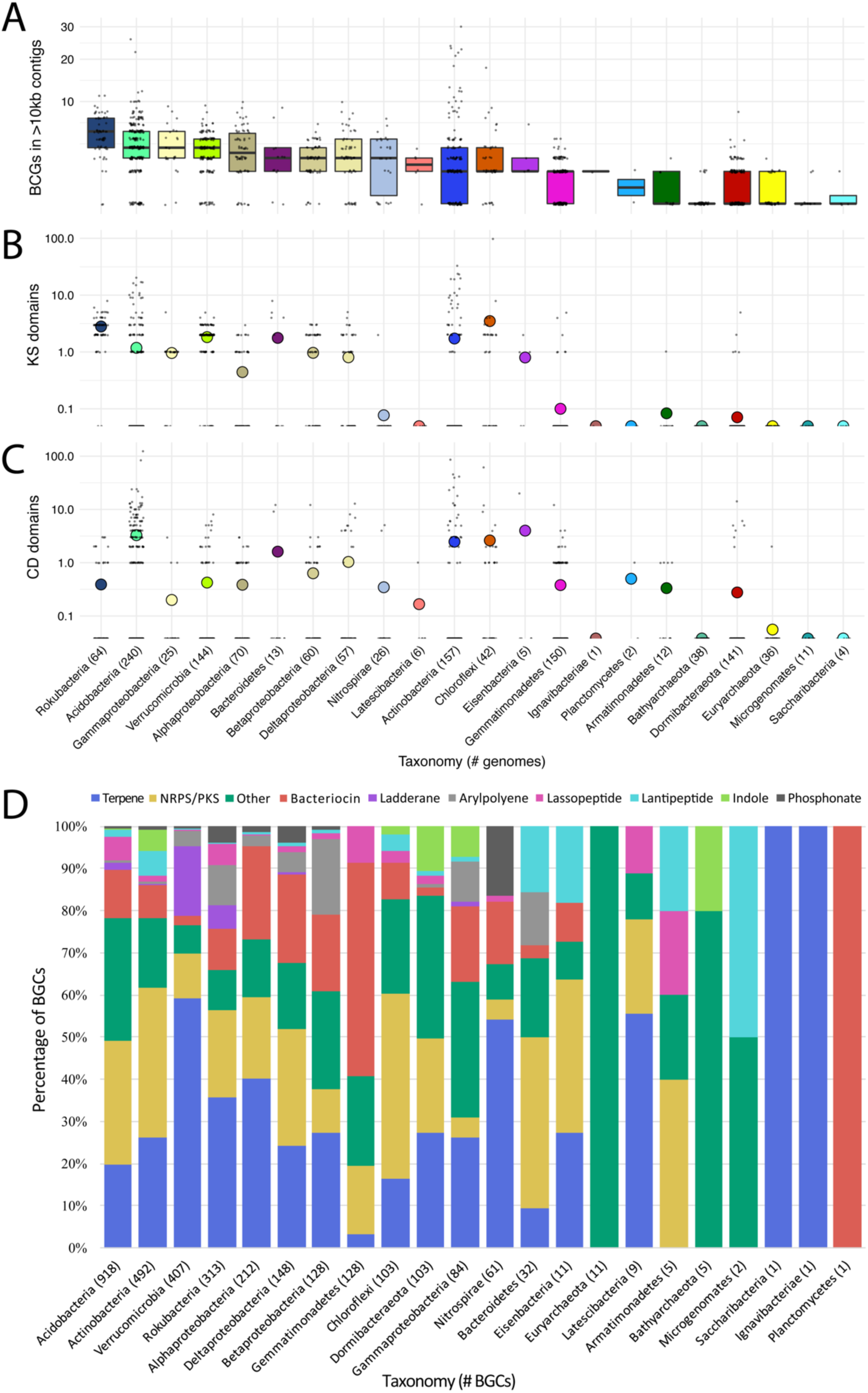
Biosynthetic gene clusters (BGCs) and key domains across taxonomic groups. **A**, BGCs per genome on contigs >10 kb, as identified by antiSMASH (square root scale). Taxonomic groups are ordered by decreasing median value (line within boxplot). Number of genomes per group in parentheses. **B**, KS domains and **C**, CD domains per genome, as called by antiSMASH (log_10_ scale). Mean group value represented as a colored dot. **D**, Percentage of BGC classes within taxonomic groups, as called by antiSMASH. Known types with <1% overall abundance grouped into ‘Other’. Total number of BGCs per group in parentheses.

The genome with the most BGCs (30) was Actinobacterium Streptosporangiales_ANG_2, found at a depth of 40 cm in hilly grassland soil (Fig. 3). A Chloroflexi genome from 20 cm meadow grassland soil (Ktedonobacter_ANG_387) had the most KS domains (99). The genome with the most CD domains (124) was a previously reported Acidobacteria from 20 cm meadow grassland soil (Eelbacter_gp4_AA13^16^). All genomes with unusually high numbers of BGCs (>15) or KS domains (>10) were classified as Actinobacteria, Acidobacteria, or Chloroflexi. Acidobacteria, Actinobacteria, Chloroflexi, Gemmatimonadetes, Deltaproteobacteria, Betaproteobacteria, Bacteroidetes and the candidate phyla Eisenbacteria and Dormibacteraeota had members whose BGCs contained >10 CD domains (Table S2).

The most common classes of BGCs identified in this study were terpenes, NRPS/PKS clusters, and bacteriocins. The classes of BGCs present within genomes depended on the taxonomic group. While ladderanes, arylpolyenes, lassopeptides, lantipeptides, indoles, and phosphonates were less common overall, some phyla had higher proportions of certain BGC classes (Fig. 4D). For example, bacteriocins are particularly prominent in genomes of Gemmatimonadetes, ladderanes were well represented among clusters in Verrucomicrobia, and phosphonates were most abundant in Nitrospirae.

Several genomes from new clades in the Actinobacteria possessed a high number of large NRPS and PKS gene clusters (Fig. S2). Those genomes with recovered 16S rRNA genes indicated that they were novel at least at the species level (Table S4). Five genomes of order Micromonosporales were recovered from both grasslands in the study, and two genomes of order Pseudocardiales were found in the Garry oak samples. The Micromonosporales and Pseudocardiales genomes encode an array of impressively large BGCs with little similarity to known BGCs from Actinobacteria in the MiBIG database (Fig. S2).

Novel clades basal to the extended class Actinobacteria were also recovered, two of which had members with significant numbers of KS and CD domains. These genomes are large (7-8 Mbp), with high GC content (>70%), and were divergent from existing publicly available Actinobacterial genomes. The largest of these clades, likely a novel family in the Streptosporangiales order, contained the genome with the most BGCs (30) found in the entire study (Streptosporangiales_ANG_2) (Fig. 3).

Three BGCs were identified within CPR genomes. This is surprising, as to our knowledge, clusters have rarely been reported in CPR genomes^56^. One CPR genome, Microgenomates_ANG_785, encodes a biosynthetic gene cluster detected as a linear azole/azoline-containing peptide (LAP). LAPs are part of the ribosomally synthesized and posttranslationally modified peptide (RiPP) natural products and are defined by the ribosomal synthesis of a precursor peptide and its subsequent post-translational modifications (PTMs)^57^. The gene cluster encodes three PTM enzyme genes that are annotated as YcaO, nitroreductase and peptidase (Fig. S3). A putative peptide precursor of 84 amino acids is present next to the YcaO gene. The YcaO acts as a cyclodehydratase that modifies the Ser and Cys residues present in the core peptide region to azolines, which are subsequently oxidized to azoles by the flavin mononucleotide-dependent dehydrogenase encoded by the nitroreductase gene^58,59^. Finally, the peptidase is proposed to cleave the leader peptide region of the precursor peptide and release the natural product. We searched for similar LAP clusters in CPR by screening a dataset enriched in CPR genomes^60^. We found 110 LAP clusters in 93 additional CPR genomes (31 Microgenomates, 51 Parcubacteria, 2 Peregrinibacteria and 1 Katanobacteria), indicating that LAPs are more widespread than previously known in CPR. Most of them (89 LAPs) contain a nitroreductase gene next to the YcaO gene. A similar LAP cluster from a soil *Rhizobium* species was recently shown to target bacterial ribosomes with highly species-specific antibiotic activity^61^.

### Biosynthetic potential with depth and vegetation

Because depth was an important factor in microbial community composition (Fig. 2), biosynthetic potential with depth was investigated. Out of 1,334 organisms, 320 significantly increased with depth (were deep enriched) and 343 significantly decreased with depth (were shallow enriched), according to DESeq2^52^ (FDR < 0.05). Most taxonomic groups had members that were deep enriched, shallow enriched, and that did not vary in abundance with depth (Fig. 5A). A few groups, such as Archaea and Nitrospirae, were primarily deep enriched whereas Gammaproteobacteria were primarily shallow enriched.

**Figure 5.**
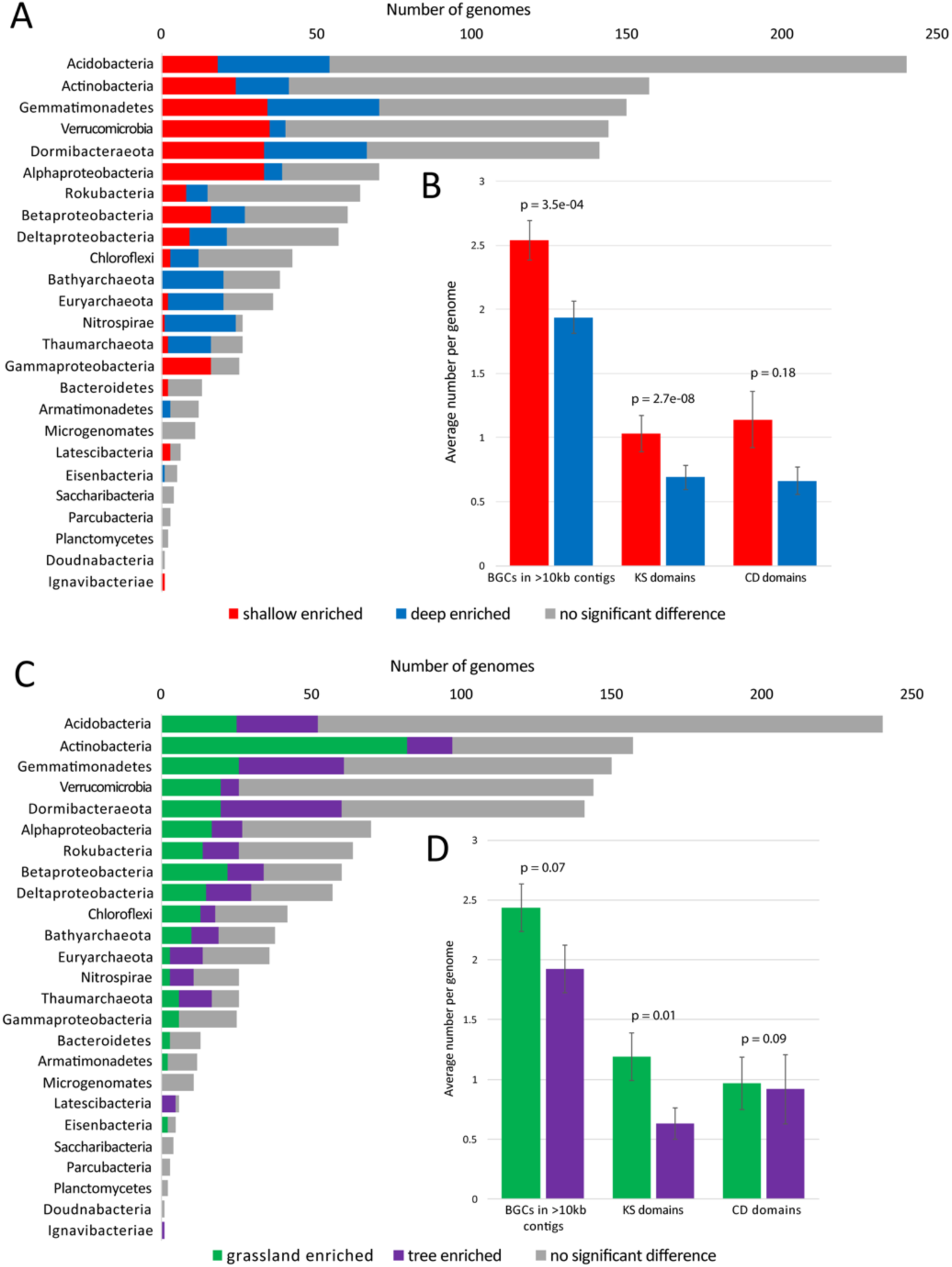
Genome, biosynthetic gene cluster (BGC), and key domain abundance with depth and vegetation. All p-values are from Wilcoxon signed-rank tests and error bars represent standard error. **A**, Number of genomes per taxonomic group that were enriched in grassland environments (green), enriched in tree-covered environments (purple), or showed no significant change (grey) with vegetation, as determined by DESeq analysis of cross-mapped, de-replicated genomes (*n* = 1,334). **B**, Average numbers of BGCs, KS domains, and CD domains per genome. **C**, Number of genomes per taxonomic group that were deep enriched (blue), shallow enriched (red), or showed no significant change (grey) with depth, determined like above. **D**, Average numbers of BGCs, KS domains, and CD domains per genome.

Average numbers of BGCs, KS domains, and CD domains per genome were compared for genomes that were deep or shallow enriched. On average, genomes of organisms enriched in shallow samples encoded more BGCs and KS domains than genomes of organisms enriched in deep samples (Fig. 5B). The overall classes of BGCs present in deep or shallow enriched genomes was not very different (Fig. S4).

In addition to depth, overlying vegetation type was tested for its significance in phylum selection and biosynthetic potential (Fig. 5C). According to DESeq2 analysis, 399 genomes were significantly enriched in grassland samples and 298 were significantly enriched in tree-covered samples. Overall, the classes of BGCs present in genomes across vegetation classes was not very different (Fig. S4). However, on average, genomes that were enriched in grasslands encoded more KS domains than genomes that were enriched at tree-covered sites (Fig. 5D).

When targeting members of particular phyla with high biosynthetic potential, it is important to consider how their abundance varies with environmental variables. Some of the genomes with at least 15 KS plus CD domains are extremely abundant at certain sampling sites or soil depths and completely absent from others (Fig. 6). The Acidobacteria with the highest biosynthetic potential were generally prevalent in meadow grassland soil. Actinobacteria were more often enriched in grassland relative to tree-covered soils, and the subset with the highest biosynthetic potential were generally prevalent in hilly grassland soil. Within the Chloroflexi, Ktedonobacter_ANG_387 was more abundant in meadow grassland soil whereas Ktedonobacter_ANG_12 was more abundant at the other sites.

**Figure 6.**
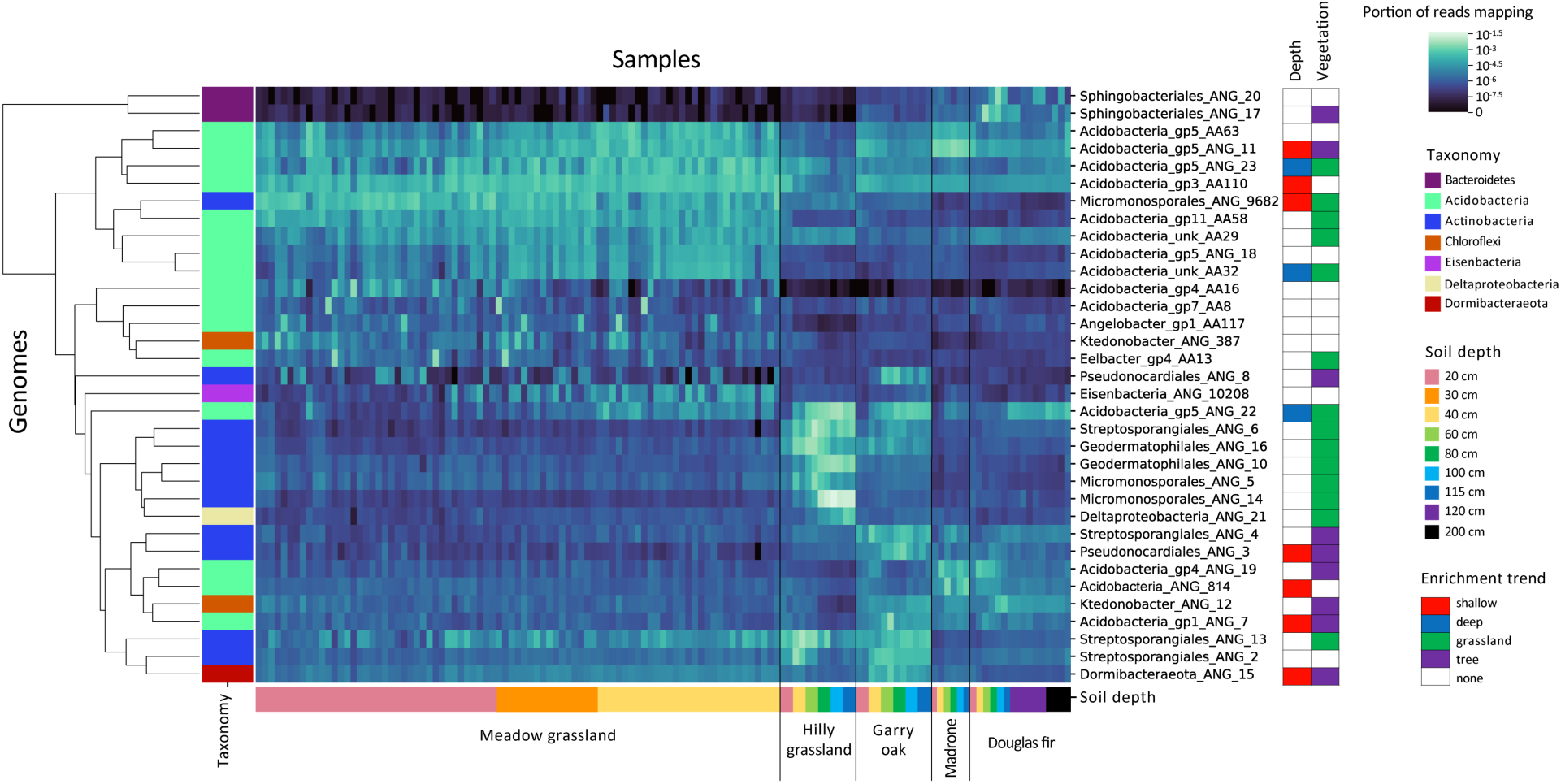
Abundance of genomes with highest biosynthetic potential across sampling sites and environments. All genomes from the high-quality, de-replicated set (*n* = 1,334) with at least 15 total biosynthetic domains were included. Lighter heatmap color, shown in log scale, indicates a higher portion of the reads in a sample (columns) mapping to a genome (rows). Genome rows were clustered based similar abundance patterns. Genome taxonomy is shown in the left vertical colored bar. Samples are grouped by site and vegetation (black vertical lines), then by soil depth (shown in horizontal colored bar). Environmental enrichment trends for each genome, as determined by DESeq, are shown in the ‘Depth’ and ‘Vegetation’ columns on the right.

## Discussion

### Understudied phylogenetic groups with high biosynthetic potential

The Actinobacteria in this study were found to have some of the highest amounts of BGCs, KS, and CD domains in their genomes. Actinobacteria with high numbers of BGCs in this study were most often novel species within the class Actinobacteria (Fig. S2, Table S4) and were often preferentially enriched in grassland relative to tree-covered (Garry oak) soil at the same site (Fig. 6). This extends findings of a prior 16S rRNA gene amplicon sequencing study by Charlop-Powers et al.^11^ that correlated Actinomycetales abundance with high NRPS adenylation and KS domain richness in soil. Therefore, when targeting Actinobacteria and their biosynthetic products, vegetation type may be an important factor.

Another phylum exhibiting high biosynthetic potential was Chloroflexi. Chloroflexi are common in soil globally, and known for their large genomes, diverse morphologies, and complex lifestyles^62^. They have been generally understudied regarding their biosynthetic potential; however, some in tropical forest soil^63^ and marine sponges^64^ were shown to encode a few PKS domains. In this study, Ktedonobacter_ANG_387 encoded 18 BGCs, with 14 classified as some type of NRPS, PKS, or hybrid combination. This enrichment is comparable to the highest degree of BGC enrichment previously shown in a few Ktedonobacteria genomes, with more NRPS/PKS clusters than reported previously^62^. Screening of compounds produced by these Ktedonobacteria showed broad antimicrobial activity^62^.

Understudied groups with notable biosynthetic potential include the candidate phyla Rokubacteria, and individual species in the Eisenbacteria and Dormibacteraeota. Rokubacteria were previously implicated in secondary metabolite production^16^, but this function has not previously been linked to Eisenbacteria or Dormibacteraeota. Although only five unique Eisenbacteria genomes were recovered, one (Eisenbacteria_ANG_10208) encoded as many CD domains as some Acidobacteria and Actinobacteria genomes with the most CD domains. These results further emphasize that phyla not historically linked to secondary metabolite production may continue to prove to be sources of potentially pharmaceutically-relevant compounds.

### Most common BGC types and their possible functions

Although terpenes were the most abundant type of BGC overall, most of their ecological functions in bacteria remain poorly understood. It has been shown that bacteria can use some terpenes to communicate with each other and with fungi^65^. Some terpenes also have antibacterial properties^66^. Because terpenes are volatile organic compounds, they have the advantage of being able to travel through both liquid- and air-filled soil pores, making them functional in a range of soil moistures. This trait may explain why they are so prevalent in these soils and saprolites which experience large shifts in soil moisture throughout the year due to the Mediterranean climate and hydrogeologic effects^24^. The wide range of novel terpene synthases in diverse soil bacteria uncovered here remain to be characterized for their function and molecular products.

The next most abundant BGC type in this study was the combined group of NRPS, PKS, and hybrid NRPS/PKS, which typically produce compounds including antibiotics, antifungals, immunosuppressants, and iron-chelating molecules^8^. After NRPS/PKS clusters, bacteriocins, which inhibit the growth of other microbes, were most prevalent overall. Bacteriocins are generally active against relatively closely-related species and likely function in reducing competition in the same niche^66^.

### Expanded phylogenetic ranges for some BGCs and possible functions

An interesting observation was the presence of clusters implicated in production of ladderanes in Verrucomicrobia. Ladderanes are only known to be produced as components of the anammoxosome membranes of anammox bacteria^67^. Anammox capabilities are only known to be present in Planctomycetes, which are part of the PVC superphylum with Verrucomicrobia^68^. The ladderanes uncovered here may serve unique, unknown functions.

We also recovered several novel BGCs for RiPPs such as lassopeptides and lantipeptides. Lassopeptide BGCs were newly found in Latescibacteria and Armatimonadetes genomes. Lassopeptides can have antimicrobial, enzyme inhibitory, and receptor antagonistic activities^69^. Further, lantipeptides are known to be widespread phylogenetically^70^, but this is the first time a cluster has been reported in a CPR bacteria genome. As lantipeptides can include lantibiotics, the finding is notable given that metabolic reconstructions for CPR bacteria consistently predict them to be symbionts^71^.

Indoles have many functions, including disruption of quorum sensing and virulence capabilities of plant pathogens and control of plant growth and root development^66^. In this study, high proportions of indole BGCs were found in Gammaproteobacteria, and in some genomes of bacteria from the newly named candidate phylum Dormibacteraeota. Interestingly, one indole BGC was also found encoded in a Bathyarchaeota genome.

Phosphonates are known to be widespread among microbes, as some have been found in Archaea^72^. While none of the few Archaeal BGCs in this study were classified as phosphonates, phosphonate BGCs were particularly abundant in Nitrospirae, which like Archaea, typically increase in relative abundance with soil depth. Phosphonates are known to function as antibacterials, antivirals, and herbicides. They also provide a mechanism to store phosphorus, which can sometimes be scarce and limiting^72^. Phosphonate use may be an adaptation of the Nitrospirae for survival in deep soil and saprolite.

### Biosynthetic capacity varies with depth and vegetation

Our finding that bacteria in shallow soil have on average higher biosynthetic capabilities than bacteria in deep soil may be attributed to the greater opportunities for interaction and competition in shallow soils, where microbial biomass and diversity are higher^73,74^. We also found that biosynthetic potential varies with vegetation type within a local environment. Some secondary metabolites, such as plant growth hormones and certain antibiotics, are produced by bacteria to benefit specific plants in their environment^75^. Previously, it was demonstrated that biosynthetic potential of amplified KS domains varies with vegetation on the continental scale^13^, and here we demonstrate similar patterns on a local scale without PCR biases.

Abundances of the different classes of BGCs were relatively consistent across genomes differentially enriched by either depth or vegetation. Few studies exist comparing biosynthetic potential across environments. However, one recent study similarly found that bacteriocin, NRPS/PKS and terpene clusters were the most common classes in 30 genomes of soil bacteria from different environments^66^. These findings suggest that while the distribution of broad classes of BGCs is mostly consistent across soil environments, the amounts of PKS and NRPS gene clusters may be environment dependent.

## Conclusion

Genome-resolved metagenomics of environmental samples allows for the discovery of new biosynthetic gene clusters and determination of the organisms and ecosystems that they reside in. Here, we uncovered environmental controls on the distribution of biosynthetic gene clusters associated with bacteria that vary in abundance with soil depth and vegetation type. This information will be useful for natural products researchers who wish to clone, isolate, or sequence the genes of these clusters. Notably, we broaden the range of phylogenetic targets for microbial products of interest, especially of NRPs and PKs. Microbial products have obvious utility in medicine and biotechnology, but they are also important for their effects on microbial communities and biogeochemical cycles. There remains much to discover about the nature of diverse secondary metabolisms in the environment.

## Acknowledgements

The authors would like to thank Sue Spalding for assistance with fieldwork and Jesse Hahm and Bill Dietrich for helpful guidance concerning the field sites. A portion of the sampling was performed at the Eel River Critical Zone Observatory, made possible by the National Science Foundation (CZP EAR-1331940). Sequencing was carried out under a Community Sequencing Project at the Joint Genome Institute. Funding was provided by the Office of Science, Office of Biological and Environmental Research, of the US Department of Energy (grant DOE-SC10010566).

## Data Availability

Sequencing reads and assembled sequences are available for 2013 and 2014 meadow grassland samples under NCBI BioProject accession numbers PRJNA297196 and PRJNA449266, respectively. Sequencing reads and assembled sequences for all other samples will be available under NCBI BioProject accession number PRJNA577476 by the time of publication. All genome sequences are available at https://doi.org/10.6084/m9.figshare.10045988.

## Supplementary Information

**Figure S1 (Separate pdf file) – High resolution version of figure 3 with genome names and bootstrap information (concatenated ribosomal protein tree of high quality, de-replicated genomes).**Black dots on branches represent bootstrap values of at least 90%.

**Figure S2.**
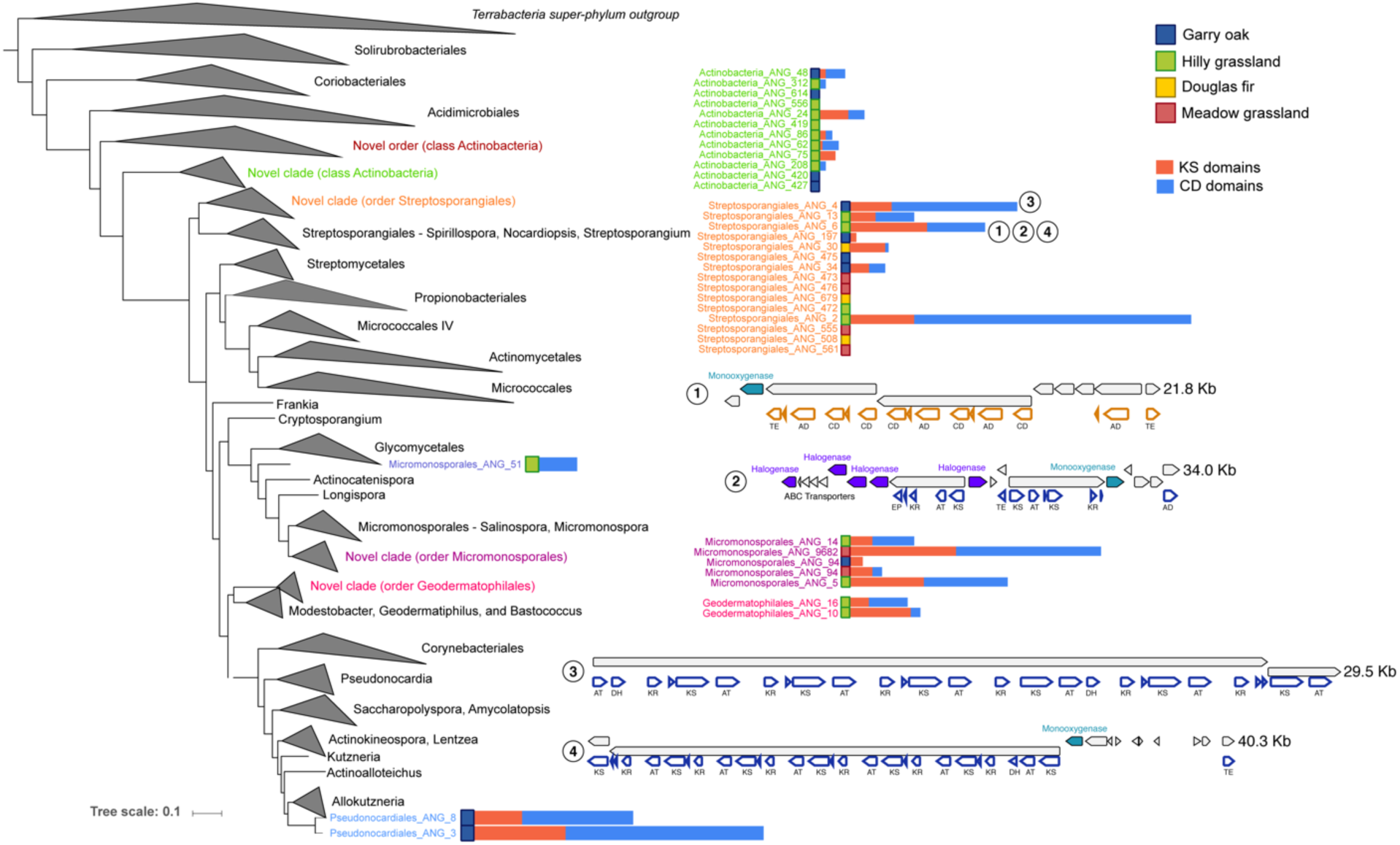
Concatenated Ribosomal protein tree of Actinobacteria. Maximum-likelihood tree based on the concatenation of 16 ribosomal proteins from all Actinobacteria genomes from this study and one reference genome from each genus on NCBI. Colored squares show which sample set the genome is from and stacked bar plots show the amount of KS (red) and CD (blue) domains identified by antiSMASH in each genome. Operon diagrams are shown for select BGCs (labelled 1-4).

**Figure S3.**
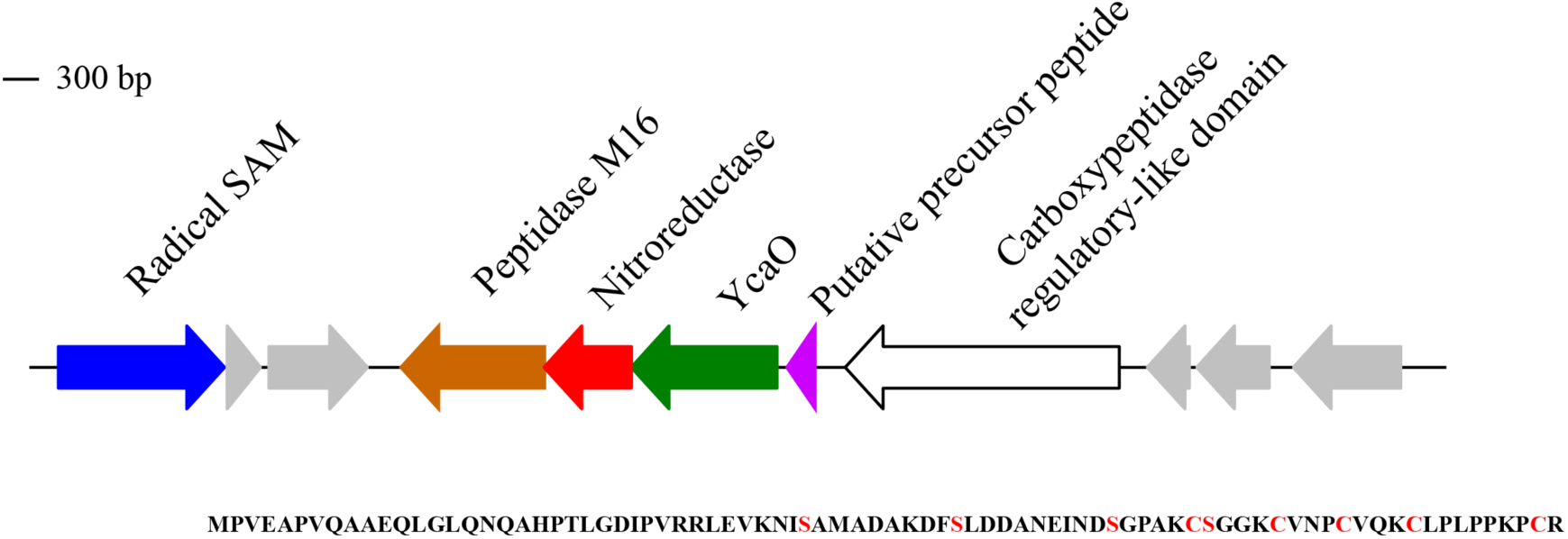
Diagram of a linear azole/azoline-containing peptide found in the candidate phyla radiation genome Microgenomates_ANG_785. The protein sequence for the putative precursor peptide is shown below with serine and cysteine residues likely modified by YcaO highlighted red.

**Figure S4.**
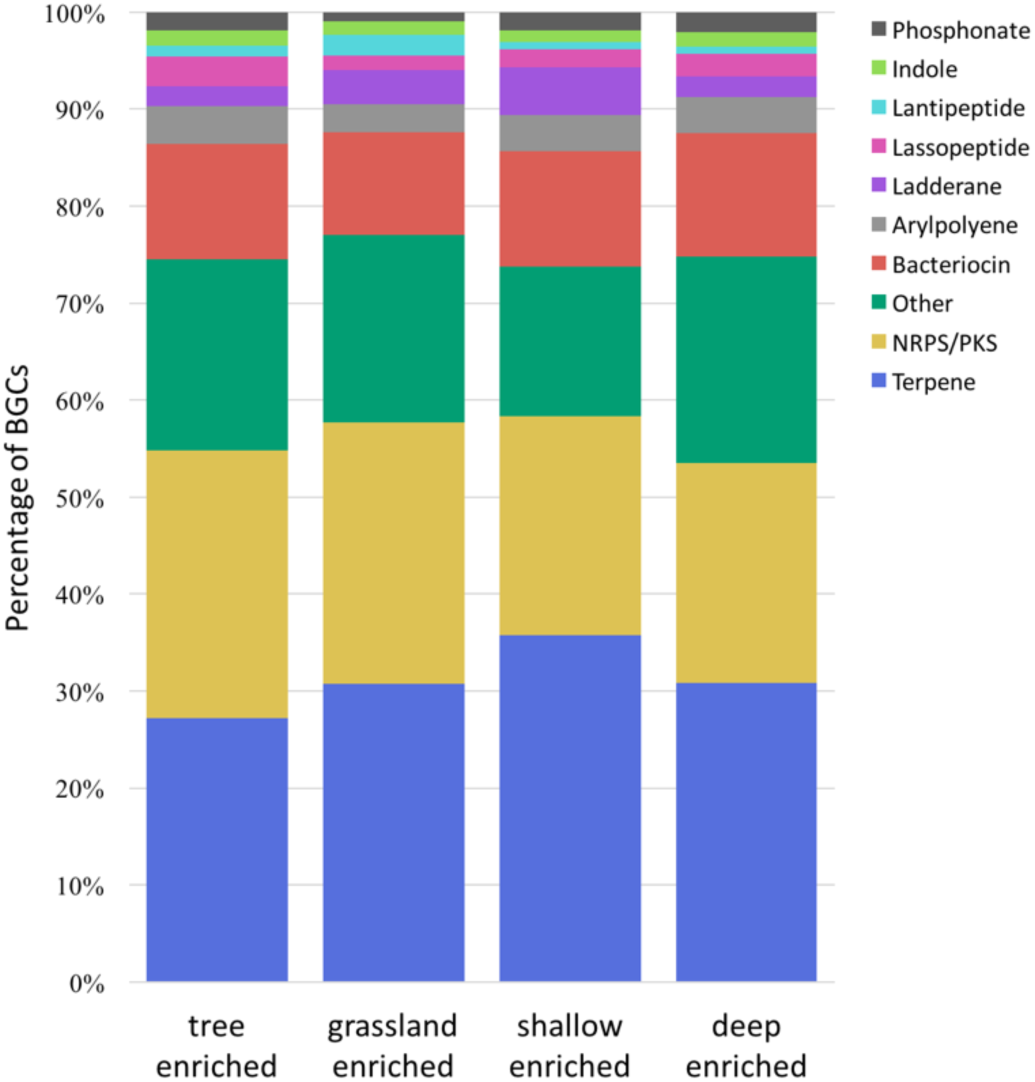
Biosynthetic gene cluster (BGC) class abundance by soil environment. Percentages of BGC classes within tree enriched, grassland enriched, deep enriched, and shallow enriched genomes. Known types with <1% overall abundance grouped into ‘Other’.

**Table S1 – Metadata for 129 metagenomic soil and saprolite samples.**

**Table S2 – Genome information, antiSMASH data, and DESeq2 results for the 1**,**334 genomes used in this study.**

**Table S3 – Biosynthetic gene clusters included in this study.**

**Table S4 – Genome information and closest 16S or ribosomal protein L6 hits of Actinobacteria genomes included in Figure S2.**

