## Supplementary figures and images for "Bacterial secondary metabolite biosynthetic potential in soil varies with phylum, depth, and vegetation type"

### Supplemental Figure S1

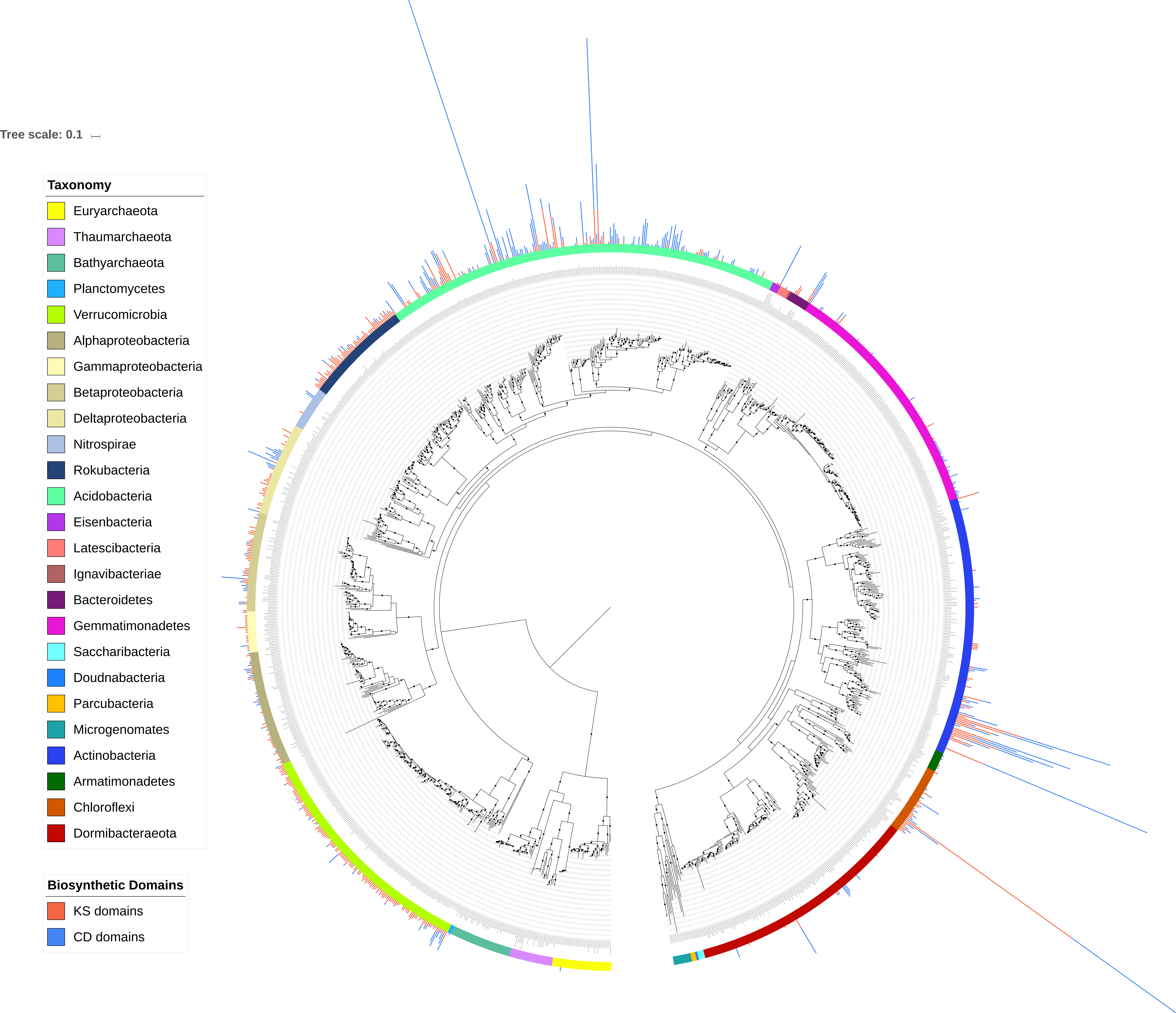
